# Scaling of G1 duration with population doubling time by a cyclin in *Saccharomyces cerevisiae*

**DOI:** 10.1101/240713

**Authors:** Heidi M. Blank, Michelle Callahan, Ioannis P.E. Pistikopoulos, Aggeliki O. Polymenis, Michael Polymenis

## Abstract

The longer cells stay in particular phases of the cell cycle, the longer it will take these cell populations to increase. However, the above qualitative description has very little predictive value, unless it can be codified mathematically. A quantitative relation that defines the population doubling time (T_d_) as a function of the time eukaryotic cells spend in specific cell cycle phases would be instrumental for estimating rates of cell proliferation and for evaluating introduced perturbations. Here, we show that in human cells the length of the G1 phase (T_G1_) regressed on T_d_ with a slope of ≈0.75, while in the yeast *Saccharomyces cerevisiae* the slope was slightly smaller, at ≈0.60. On the other hand, cell size was not strongly associated with T_d_ or T_G1_ in cell cultures that were proliferating at different rates. Furthermore, we show that levels of the yeast G1 cyclin Cln3p were positively associated with rates of cell proliferation over a broad range, at least in part through translational control mediated by a short uORF in the *CLN3* transcript. Cln3p was also necessary for the proper scaling between T_G1_ and T_d_. In contrast, yeast lacking the Whi5p transcriptional repressor maintained the scaling between T_G1_ and T_d_. These data reveal fundamental scaling relations between the duration of eukaryotic cell cycle phases and rates of cell proliferation, point to the necessary role of Cln3p in these relations in yeast and provide a mechanistic basis linking Cln3p levels to proliferation rates and the scaling of G1 with doubling time.

## INTRODUCTION

Recurring shapes and patterns in nature are sometimes described with mathematical relationships. As a result, these natural processes can be predicted and understood better. Regarding patterns of eukaryotic cell division, one could ask: How are the lengths of eukaryotic cell cycle phases related to each other and to the total doubling time of the population? Can such relations be described mathematically, in the form of a scaling formula? If so, what are the molecular mechanisms that govern the scaling? A scaling relation that describes eukaryotic cell division would be a significant advance. For example, it could serve as a point of reference against which the effects of genetic or other perturbations can be evaluated.

The ‘textbook’ view in the coordination of growth and division in the eukaryotic cell cycle (e.g., see Fig. 10-26 in (MORGAN 2007)) is that expansion of the G1 phase of the eukaryotic cell cycle accounts for most, if not all, of the lengthening of the cell cycle in slower proliferating cells, in budding yeast (JOHNSTON *et al*. 1977; BRAUER *et al*. 2008) or humans (BASERGA 1985; FISHER 2016). However, there is no report in the literature of a quantitative relation that defines the doubling time (T_d_) as a function of the time yeast or human cells spend in the G1 phase (T_G1_). Here, based on all the available data for budding yeast and human cell populations, we derived for the first time in the field scaling relations between T_G1_ and T_d_. These scaling relations also allowed us to critically evaluate the role of cell cycle regulators in yeast cells proliferating at different rates.

Two key regulators of the length of the G1 phase in *S. cerevisiae* are the Cln3p and Whi5p proteins. The G1 cyclin Cln3p promotes initiation of DNA replication (CROSS 1988; NASH *et al*. 1988). In contrast, the transcriptional repressor Whi5p acts analogously to the retinoblastoma gene product in animals, to inhibit the G1/S transition (COSTANZO *et al*. 2004; DE BRUIN *et al*. 2004; PALUMBO *et al*. 2016). It has been reported that while synthesis of Cln3p parallels cell size, the synthesis of Whi5p is independent of cell size (SCHMOLLER *et al*. 2015), arguing that dilution of Whi5p as cells get bigger in G1 governs the length of the G1 phase (SCHMOLLER AND SKOTHEIM 2015; SCHMOLLER *et al*. 2015).

Here, we obtained the first measurements of Cln3p and Whi5p levels as a function of proliferation rates in steady-state cultures. The levels of Cln3p varied over a broad range, due to a uORF affecting translation of *CLN3*. Our data also show that loss of Whi5p does not significantly affect the scaling relation between T_d_ and T_G1_. Instead, we provide strong evidence for the functional and molecular basis for the necessary role of Cln3p in this process.

## MATERIALS AND METHODS

### Strains

Unless stated otherwise, *S. cerevisiae* wild-type, *cln3Δ* and *whi5Δ strains* were in the BY4741 background (NCBI Taxon 559292; *MATa, his3Δ1, leu2Δ0, ura3Δ0, met15Δ0*) and they have been described previously (SOMA *et al*. 2014). For protein surveillance, we constructed an otherwise wild type strain that carried epitope-tagged *WHI5* and *CLN3* alleles at their endogenous chromosome locations. First, a commercially available *WHI5-TAP::HIS3* strain (BY4741 otherwise; GE Healthcare) was backcrossed three times into the W303 background (NCBI Taxon 580240; *MATa leu2-3,112 trp1-1 can1-100 ura3-1 ade2-1 his3-11,15*). Then, it was crossed with an otherwise wild-type strain carrying a *CLN3-13MYC* allele (W303 background) described elsewhere (THORBURN *et al*. 2013), and kindly provided by Dr. A. Amon (MIT and HHMI). The resulting diploid was sporulated and dissected, to obtain *MATa* haploid segregants carrying both the epitope-tagged *WHI5* and *CLN3* alleles (strains HB94/97; *MATa CLN3-13MYC::TRP WHI5-TAP::HIS leu2 ura3 met15*), which were used in the experiments shown in Figure 4. We verified expression of Whi5p-TAP and Cln3p-(Myc)_13_ in this strain (see Supplementary File 1), and their absence in *whi5Δ* or *cln3Δ* strains, respectively. We also generated a derivative of this strain, which lacks the uORF in the 5’-leader of the *CLN3* mRNA. To this end, we used plasmid A-315T-pMT10 we had described previously (POLYMENIS AND SCHMIDT 1997), as a template in a PCR reaction with forward (5’-CAAGAACTACCATTCGACAGG-3’) and reverse primers (5’-CGTACAGAAAGCGTATCAAA-3’) to generate a product that carries in the 5’-leader of *CLN3* the *URA3*-marked A-315T mutation that inactivates the uORF. We then used this PCR product to transform strain HB94 (*WHI5*-*TAP, CLN3-13MYC*). Genomic DNA of transformants was sequenced to verify the presence of the A-315T mutation. Confirmed A-315T mutants were then backcrossed with wild type (W303) to segregate away possible secondary mutations at other loci. The resulting heterozygote was sporulated and dissected to isolate a *WHI5-TAP, A-315T-CLN3-13MYC* segregant (HB104), which was used in the experiments shown in Figure 4.

### Datasets for population-based cell cycle parameters

All the obtained variables we report here represent population averages. They do not resolve intergenerational differences in cell cycle progression of the same cells in successive cell cycles. In the context of this study, population averages hold significant advantages: First, they are easily obtained; Second, they are ubiquitously used and reported in the literature; Third, they allow straightforward comparisons between different systems, for example between yeast and human cells (see Figure 1).

**Figure 1.**
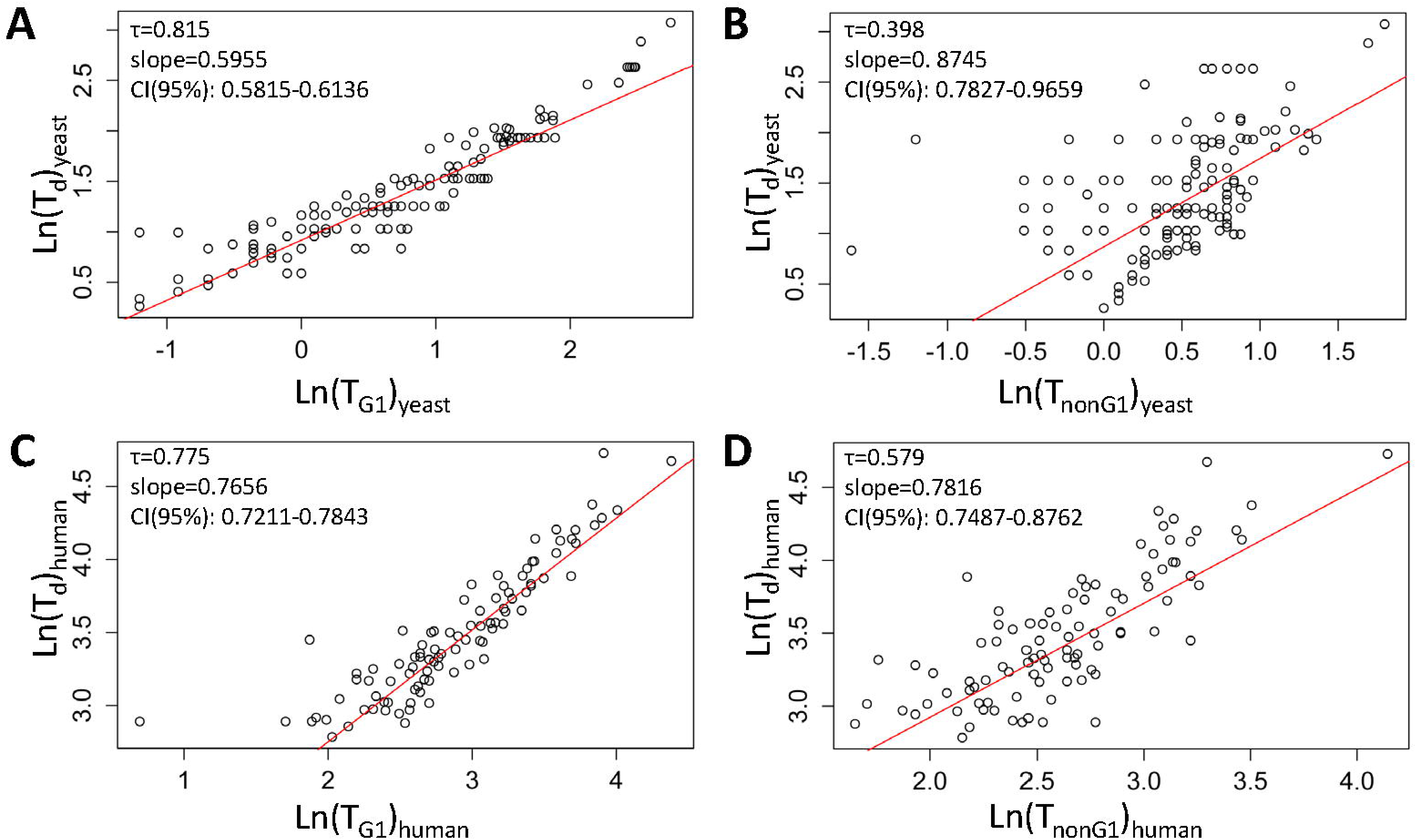
Linking the length of the G1 phase with population doubling time. Scatter plots of T_G1_ (**A, C**) or T_nonG1_ (**B, D**) values on the x-axis, against T_d_ values (y-axis). All plots used the natural logarithms of the values for yeast (**A, B**) and human (**C, D**) cells from Tables S1 and S2, respectively. The Kendall’s (τ) rank correlation coefficient is shown in each case. In red are regression lines of the Siegel repeated medians. The slope and the associated 95% confidence intervals of the linear model are shown in each case. Additional statistical parameters associated with these plots are shown in Table 1.

**Figure 2.**
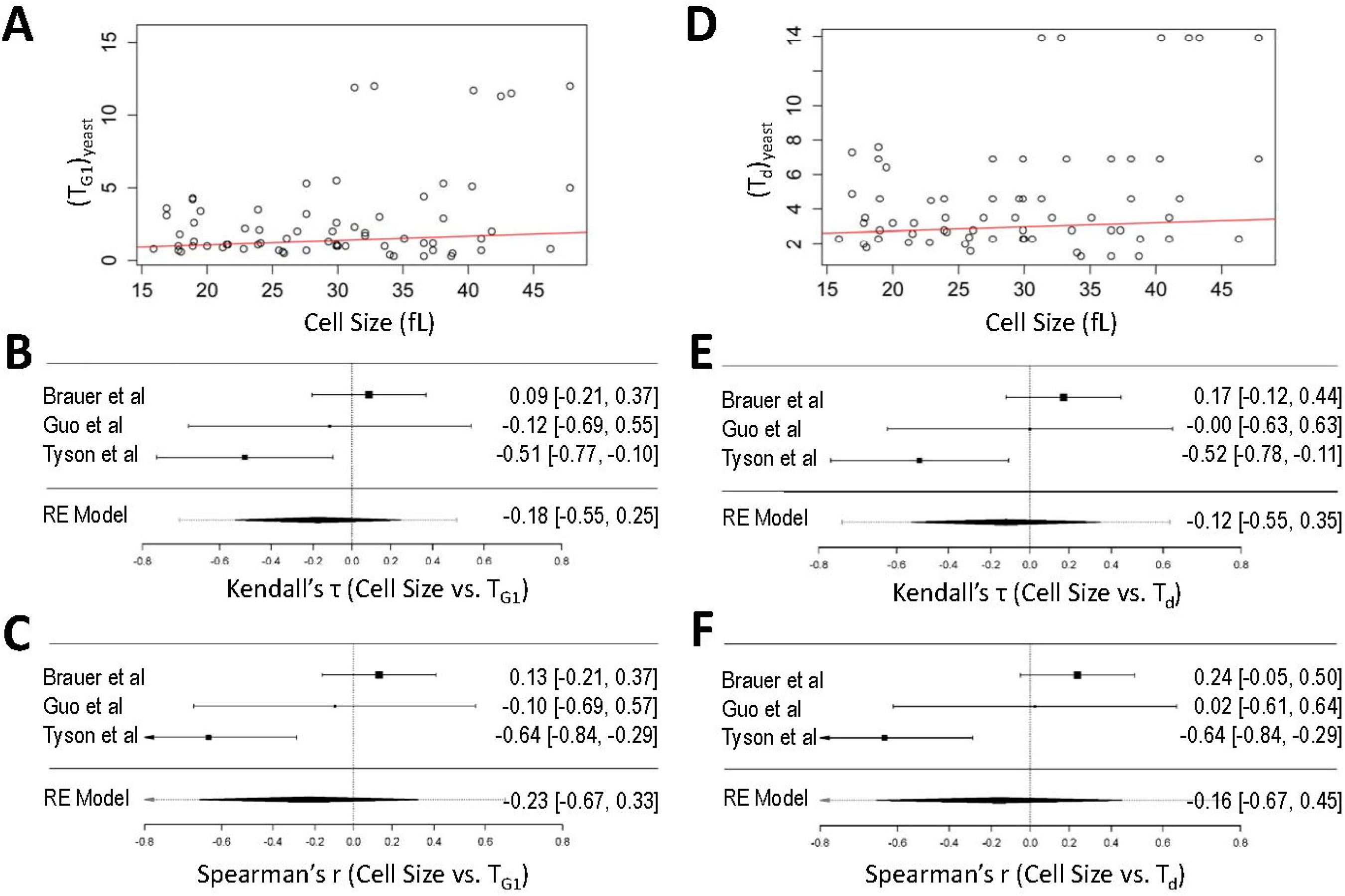
Cell size does not correlate with T_d_ or T_G1_ in yeast. Scatter plots of cell size values (x-axis) against T_d_ (**A**) or T_G1_ (**D**) values (y-axis) from the data shown in Table S1. In red are regression lines of the Siegel repeated medians. Forest plots of the measure of effect for each of the studies included in the analysis (TYSON *et al*. 1979; GUO *et al*. 2004; BRAUER *et al*. 2008), based on the Kendall’s (τ) rank correlation coefficients (**B, E**), or Spearman’s (r) rank correlation coefficients (**C, F**), are shown in each case, for cell size vs. T_G1_ (**B, C**) and cell size vs. T_d_ (**E, F**). The confidence intervals from each study are shown in parentheses and represented by horizontal whisker lines. In the studies in which the confidence intervals overlap with the vertical line at the 0 point on the x-axis, their effect sizes do not differ from no effect. The meta-analyzed measure of the effect is shown at the bottom of each plot, based on random effects (RE) models.

For yeast, the data we collected (Table S1) were from wild type strains from various backgrounds, except in the few cases where they carried temperature-sensitive alleles, such as *cdc* mutations (JAGADISH AND CARTER 1977), to estimate the length of the G1 phase upon transfer to the non-permissive temperature. The methods used to calculate the fraction of G1 cells included: measurements of the DNA content of the cells by flow cytometry (SLATER *et al*. 1977; JOHNSTON *et al*. 1980; GUO *et al*. 2004; BRAUER *et al*. 2008; HENRY *et al*. 2010); budding (TYSON *et al*. 1979; RIVIN AND FANGMAN 1980); sensitivity to cell cycle arrest before DNA replication by pheromone (HARTWELL AND UNGER 1977; JAGADISH AND CARTER 1977), or *cdc* (JAGADISH AND CARTER 1977) mutations. In this study, to obtain the fraction of G1 cells (e.g., see Figures 3, 4), we used DNA content measurements by flow cytometry, as described previously (HOOSE *et al*. 2012; HOOSE *et al*. 2013).

**Figure 3.**
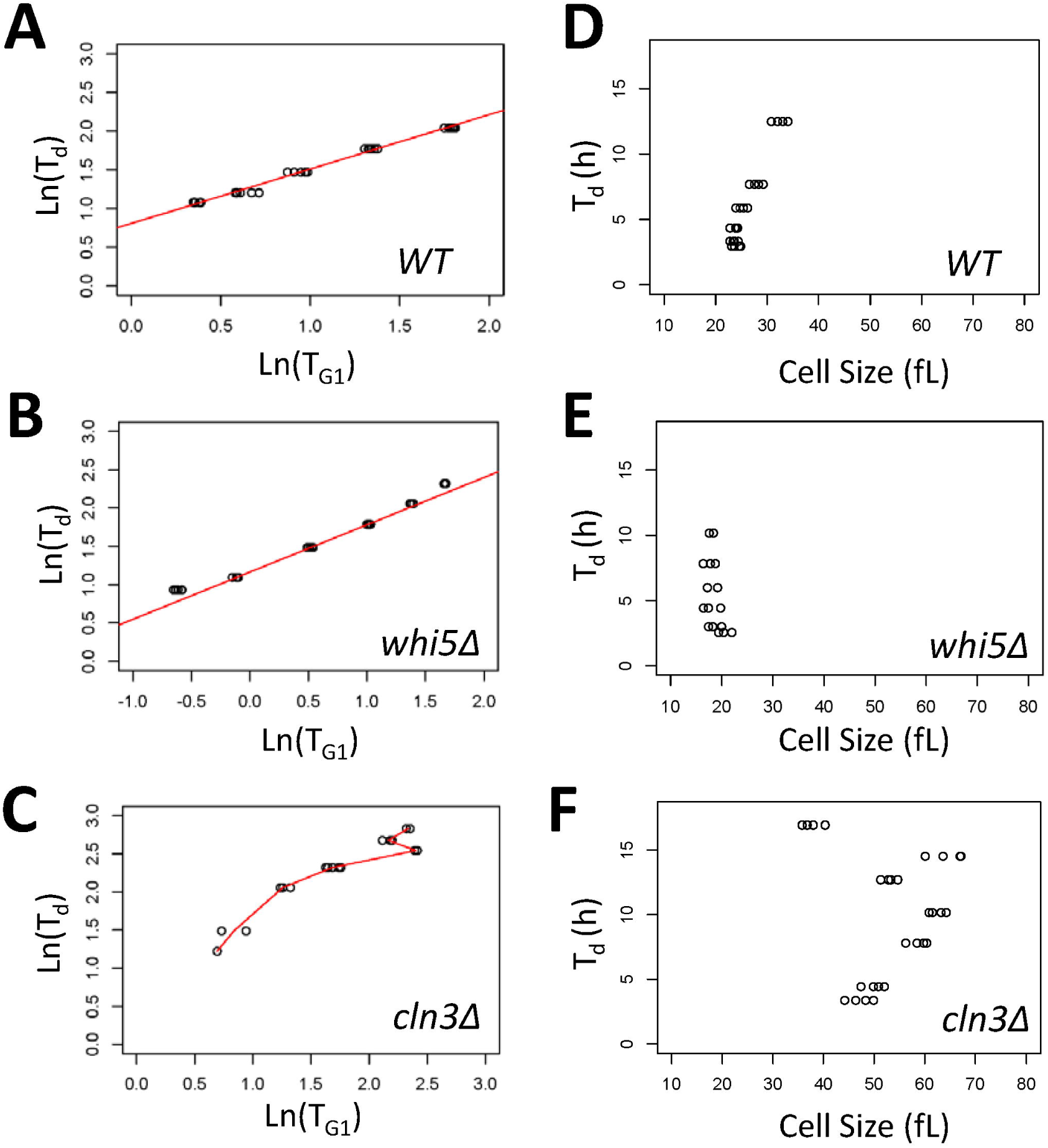
Cln3p, but not Whi5p, imposes the proper relation between T_G1_ and T_d_. Scatter plots of T_G1_ values on the x-axis, against T_d_ values (y-axis). All plots used the natural logarithms of the values for wild type (**A**) whi5Δ (**B**) or cln3Δ (**C**) cells, sampled from chemostat cultures several times at each dilution rate, as indicated. For wild type and whi5Δ cells, in red are regression lines of the Siegel repeated medians, and the slope of the linear models are shown (additional statistical parameters are in Table 1). For cln3Δ cells, the red line shown simply connects the average values at each dilution rate. There is no regression line because the relation between T_d_ and T_G1_ breaks down, especially at longer generation times. Scatter plots of the relation of cell size and T_d_ in wild type (**D**) or cells lacking Whip5 (**E**) or Cln3p (**F**), with cell size values (x-axis) plotted against T_d_ (y-axis) from the same cultures described in (**A-C**). All the strains were in the BY4741 background (see Materials and Methods).

**Figure 4.**
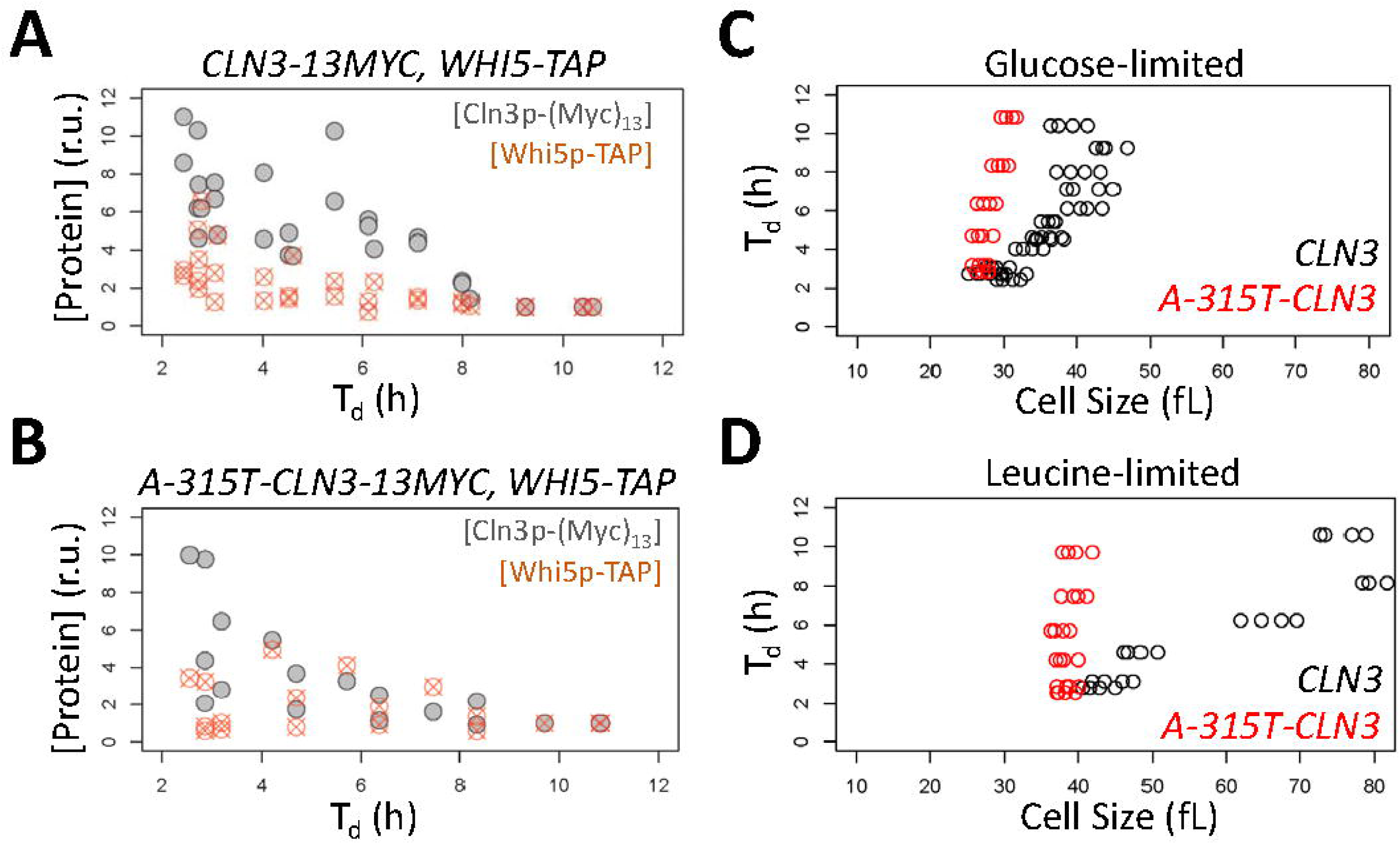
The levels of the G1 cyclin Cln3p vary over a broad range as a function of T_d_, due to a uORF affecting translation of CLN3. Scatter plots of the relative abundance (y-axis) of Cln3p-(Myc)_13_ and Whi5p-TAP in otherwise wild type CLN3-13MYC, WHI5-TAP cells (**A**), or CLN3 uORF (A-315T-CLN3) mutant cells (**B**), against T_d_ (x-axis). Each data point in the scatter plots is the average of immunoblot signal intensities run in duplicate, and detected with antibodies against the Myc or TAP epitopes (see Materials and Methods). All the raw immunoblots used to quantify protein levels are shown in the Source Data (File S1). Before averaging, each individual signal intensity value was normalized against loading in the corresponding immunoblot lane (visualized with Ponceau staining, see File S1 and Materials and Methods). For each Cln3p-(Myc)_13_ or Whi5p-TAP relative unit (r.u.) shown in the scatterplots (**A, B**), the normalized, averaged intensities were scaled by the lowest value (set to 1) for each protein in the given chemostat experiment run at different dilution rates. Scatter plots of the relation of cell size (x-axis) and T_d_ (y-axis) in the indicated strains, cultured under glucose (**C**) or leucine (**D**) limitation, from the same cultures described in (**A**) and (**B**).

For human cells, earlier studies employed ^3^H-thymidine pulses or division waves after thymidine block (BASERGA 1985). The doubling times of the NCI-60 human cancer cell lines we included in Table S2 are known (ROSS *et al*. 2000; SCHERF *et al*. 2000; POLYMENIS 2017), but there was no quantitative cell cycle data for most of the cell lines. However, images of DNA content profiles for the NCI-60 panel, albeit with no quantification, have been published (GARNER AND EASTMAN 2011). We requested and obtained high-resolution files of these images from Dr. Alan Eastman (Geisel School of Medicine - Dartmouth College).

From the entire DNA content histogram, to quantify the fraction of cells in G1, we used imaging software to measure the area on the left side of the G1 peak (from peak to valley) and multiplied this area by two, as has been described previously (JOHNSTON *et al*. 1980). This approach avoids complications from heavy right side tails due to S phase cells and yields an acceptable estimate of the relative G1 length as a fraction of total cell cycle time. We then combined these values with all others available from human cells, cancerous and normal (SISKEN AND KINOSITA 1961; DEFENDI AND MANSON 1963; LENNARTZ AND MAURER 1964; AOKI AND MOORE 1970; BASERGA 1985; KUMEI *et al*. 1989; BRONS *et al*. 1992; LUCIANI *et al*. 2001; HAHN *et al*. 2009), to compile a dataset of 96 values for G1 length (T_G1_) and doubling time (T_d_) for human cells, shown in Table S2.

### Estimates of G1 length

The values we show in Tables S1 and S2 were obtained from studies reporting on the *relative* duration of the G1 phase. To estimate the *absolute* length of the G1 phase, T_G1_, we multiplied the relative G1 length by T_d_ (Tables S1, S2). This simple equation is the appropriate one to use with chemostat data when as many cells are removed as are being produced in the culture (HOFFMAN 1949). Subtracting T_G1_ from T_d_ yields the duration of the rest of the cell cycle phases (T_nonG1_). For non-chemostat data, other more elaborate equations could be used, especially for the asymmetric patterns of division of budding yeast (HARTWELL AND UNGER 1977; JOHNSTON *et al*. 1980). However, for simplicity and ease of comparison across systems, we uniformly applied the simple equation mentioned above. Furthermore, inaccuracies in the absolute values of T_G1_ may only affect the intercept of the linear relation between T_d_ and T_G1_, but not the slope that describes the fundamental scaling between T_d_ and T_G1_, or any of our conclusions. Indeed, when we plotted in the same manner as in Figure 1 only the chemostat values in Table S1 (94 data points), the slope of LnT_d_ ∝ LnT_G1_ was 0.6107. For the remaining, non-chemostat values (55 data points), the slope of LnT_d_ ∝ LnT_G1_ was 0.6074.

Lastly, a limitation of the non-chemostat data in yeast and all the data from human cells is that it is assumed that cell death contributes negligibly to the doubling time of the population. This assumption is reasonable in yeast because young cells vastly outnumber older ones approaching senescence. However, it may be of concern in mammalian culture systems. Hence, our data with human cells should be interpreted with caution, because the fraction of growing cells in the culture may be significantly lower than one. Also, although we are looking at trends that seemingly hold across a multitude of human cell types, the data were overwhelmingly derived from cancer cell lines, which in many cases have altered cell cycles.

### Chemostat cultures

The experiments were done using a New Brunswick BioFlo (BF-110) reactor with a working volume of 880 mL. The reactor was run at room temperature, as described earlier (HENRY *et al*. 2010). In each experiment and at each dilution rate the reactor was sampled several times to measure protein levels by immunoblots, the DNA content with flow cytometry, and the cell size and cell density of the culture using a Beckman Z2 channelyzer (HENRY *et al*. 2010; HOOSE *et al*. 2012), as indicated. We measured the cell density at every sampling, to ensure that we never reached ‘wash-out’ conditions at the high dilution rates. In every experiment the cell density remained >1E+07 cells/ml and did not vary more than 3-fold between the lowest and highest dilution rates.

### Protein surveillance

Proteins were resolved onto 4-12% Tris-Glycine gels (Thermo Scientific, Cat#: XP04125BOX). Cln3p-(Myc)_13_ was detected with an anti-Myc antibody (Abcam, Cat #: ab13836). All other procedures for TAP-tagged protein detection, extract preparation for immunoblots and their analysis have been described elsewhere (BLANK *et al*. 2017).

### Statistical analysis

Data were analyzed and displayed with R language packages. All R functions, the corresponding packages, and their use are listed in Table S3. To build the linear models we described using the values for the yeast (Table S1) and human (Table S2) datasets, we first examined if the assumptions for building simple, linear parametric models were satisfied. The diagnostic residual plots evaluating whether the errors were independent of each other, normally distributed around a mean of zero and equal variance, are shown in Figure S2. In both the yeast and human datasets, the existence of a few outlier points appeared to violate the necessary assumptions (Figure S2; p<0.05 for assessment of the assumptions using the global test on four degrees-of-freedom (PENA AND SLATE 2006)). Hence, we opted for non-parametric, robust linear regression models based on Siegel repeated medians (Table 1). For the meta-analysis of cell size data (see Figure 2) we used the *metafor* R language package (VIECHTBAUER 2010). Briefly, the correlation coefficients from each study were transformed using Fisher’s z transformation. An unbiased random-effects analysis, as opposed to a fixed-effects one, was then performed using this index, and the summary values were converted back to correlations and displayed as such with forest plots (Figure 2, Table S3).

### Data availability

Strains and plasmids are available upon request. The authors affirm that all data necessary for confirming the conclusions of the article are present within the article, figures, and tables. Figure S1 shows goodness-of-fit plots for lognormal distribution of yeast and human T_G1_ values. Figure S2 shows diagnostic plots of simple linear regression models for T_d_ and T_G1_ values of yeast and human cells. Figure S3 shows additional chemostat experiments with *whi5Δ* and *cln3Δ* cells. Table S1 lists all the cell cycle values for yeast cells from the literature. Table S2 lists all the cell cycle values for human cells from the literature. Table S3 lists all R functions, the corresponding packages, and their use. File S1 contains all raw immunoblot images generated and used in this study. All supplementary figures, tables and files can be found at (10.6084/m9.figshare.7011275).

## RESULTS

### Rationale

The rationale for the experiments we describe was the following: First, use all the available values from the literature to derive a quantitative relationship of the population doubling time (T_d_) as a function of the time eukaryotic cells spend in the G1 phase of the cell cycle (T_G1_) (Figure 1). Second, based on the same datasets and analyses, examine if cell size is also related to T_d_ or T_G1_, because cell size is often used as a proxy for the control of cell division by nutrients (Figure 2). Third, use the linear relation linking T_d_ and T_G1_ as a metric to evaluate the contributions of Whi5p and Cln3p, two proteins that govern the G1/S transition in budding yeast (Figure 3). Fourth, if Cln3p or Whip5p impinges on the relation between T_d_ and T_G1_, then provide a mechanistic understanding of its role (Figure 4).

### T_G1_ values are distributed lognormally, consistent with exponential patterns of growth

We compiled the available values for T_d_ and T_G1_ from the literature for budding yeast (Table S1) and human (Table S2) cells (see Materials and Methods). With the dataset of T_G1_ values at hand, we next examined their distribution. Knowing how the T_G1_ values are distributed will inform how to better model T_G1_ against T_d_ and offer some insight into the processes that determine G1 length. We found that T_G1_ values were not normally distributed for yeast (p = 4.434E-14, Shapiro-Wilk test) or human cells (p = 1.039E-07, Shapiro-Wilk test). Instead, T_G1_ values fit better a lognormal distribution. For example, for yeast T_G1_ values, the Anderson-Darling statistic was the lowest for the lognormal distribution (0.347), compared to other distributions (Weibull: 1.693; gamma: 1.521; exponential: 2.462). As expected for lognormal distributions, log-transformed values of T_G1_ were normally distributed for yeast (Figure S1A-D, p = 0.1871, Shapiro-Wilk test) and human cells (Figure S1E-H, p = 0.3099, Shapiro-Wilk test). The apparent lognormal distribution of T_G1_ values is consistent with a multiplicative process of many, positive, independent random variables that determine the G1 length (KOCH 1966). Lognormal distributions are very common in biological growth processes (MOSIMANN 1988). In cell proliferation, lognormality has been proposed to reflect exponential patterns of growth in mass. Despite fluctuations in the growth rate constant, the growth of the overwhelming majority of cellular components is influenced similarly, leading to lognormality (KOCH AND SCHAECHTER 1962; KOCH 1966). In budding yeast and other cell types there is evidence for exponential patterns of protein synthesis (ELLIOTT AND MCLAUGHLIN 1978; DI TALIA *et al*. 2007; TZUR *et al*. 2009) and increase in mass in the cell cycle (BRYAN *et al*. 2012; SON *et al*. 2012). Such considerations accommodate the lognormality of T_G1_ values we describe here.

### Strong association between T_G1_ and T_d_, but non-G1 phases also expand in lower proliferation rates

To test for association between T_G1_ and T_d_ we used the distribution-free Spearman’s and Kendall’s tests for independence based on ranks. We used these non-parametric, distribution-free tests because of the existence of outliers even in log-transformed T_G1_ values (e.g., see Figure S1). The high values (> 0.75) of the rank correlation coefficients (τ for Kendall’s and r for Spearman’s; see Table 1) show a strong positive association between T_G1_ and T_d_, for both yeast (Figure 1A) and human (Figure 1C) cells. Interestingly, however, the duration of the non-G1 phases (T_nonG1_) of the cell cycle were also positively correlated with T_d_ (Figures 1B, D; and Table 1) in both organisms, albeit less so in yeast (τ=0.398; Figure 1B) than in human cells (τ=0.579; Figure 1D). Overall, our data document the strong association between T_G1_ and T_d_ (Figures 1A, C). Additionally, they suggest that growth requirements for cell division are not registered exclusively in G1, but also later in the cell cycle (see Figures 1B, D), in agreement with observations from other groups (ANASTASIA *et al*. 2012; FERREZUELO *et al*. 2012; DOWLING *et al*. 2014; SOIFER AND BARKAI 2014; CERULUS *et al*. 2016; MAYHEW *et al*. 2017; GARMENDIA-TORRES *et al*. 2018).

### A scaling relation between T_G1_ and T_d_

To estimate a predictive and quantitative relation between T_G1_ and T_d_ we derived non-parametric, robust linear regression models using the Siegel repeated median estimates (SIEGEL 1982), (Figure 1 and Table 1; see also Materials and Methods). The intercepts of the linear T_G1_ vs. T_d_ plots reflect the apparent minimum duration of the S+G2+M phases in yeast (1.4 h; Table 1) and human (7.6 h; Table 1) cells. The slopes in the linear relations indicate how much T_d_ is affected by T_G1_, or T_nonG1_. For example, if non-G1 phases were not expanding in slower proliferating cells, then one would expect a vertical line parallel to the y-axis in T_nonG1_ vs. T_d_ plots. We noticed that the slope of the regression of T_G1_ on T_d_ appeared to slightly differ between the yeast and human datasets (Figure 1 and Table 1). Applying the non-parametric Sen-Adichie test for parallelism confirmed that the difference in the slopes of the regression lines of T_G1_ on T_d_ between yeast and human cells was statistically significant (V statistic = 7.324, p-value = 0.007). Although the T_G1_ distributions themselves are lognormal, log-transformation is not necessary for any of our conclusions, since we used a non-parametric, ordinal-based analysis in all our statistical tests. Nonetheless, we display regression plots using log-transformed data to improve visualization, because data points appear more evenly on these graphs. Furthermore, log-transformed values are often incorporated in scaling relations between the measured variables in the literature (CHAN AND MARSHALL 2010). The quantitative relations we identified linking T_G1_ with T_d_ are significant because they provide a framework to interpret experimental perturbations in cell cycle progression and cell proliferation, as we will describe for yeast cells.

### Nutrient-specific, but not growth rate-dependent association between cell size and T_G1_, or T_d_

Control of cell size has frequently been used synonymously with growth control of G1 transit in the cell cycle, especially in budding yeast. Daughter cells of *S. cerevisiae* are born smaller than their mother is, and they will not initiate a new round of cell division until they reach a size characteristic of the culture medium. The rate at which daughter cells increase in volume has been reported to contribute to the size at which they will initiate a new round of cell division (FERREZUELO *et al*. 2012). Note that unless indicated otherwise, here we use the term ‘growth rate’ to describe the rate at which cells proliferate, and not the rate at which they increase in size in a given cell cycle. As the cell cycle is prolonged in poor nutrients, it is also widely assumed that the cells get smaller (e.g., see Fig. 10-26 in (MORGAN 2007)). To test the strength of the association between cell size and T_G1_ or T_d_, we combined the available data from previous studies ((TYSON *et al*. 1979; GUO *et al*. 2004; BRAUER *et al*. 2008); see Table S1). From such an unbiased but unweighted analysis (Figure 2A, D), it appeared that the size of yeast cells was not significantly associated with T_d_ (p-value= 0.171, based on Kendall’s test; Figure 2A) or T_G1_ (p-value= 0.2449, based on Kendall’s test; Figure 2D).

Unlike the strong association of T_G1_ with T_d_, which was consistent across studies (see Figure 1 and Materials and Methods), the association between cell size and T_G1_ or T_d_ appeared to vary among the relevant studies. Given the different number of samples analyzed in each study and their associated variance, we calculated the effect sizes from each study separately, based on the non-parametric Spearman’s and Kendall’s correlation coefficients. These study-specific correlation coefficients then served as the effect size index, to standardize the different studies and arrive at a summary correlation (BORENSTEIN 2009). The results were visualized in typical ‘forest’ plots (Figures 2B, C, E, F). A negative association between size and rates of cell proliferation, with cells getting smaller with larger T_d_ values, was only evident at a moderate level in batch cultures (τ = −0.52, r = −0.64; see Figures 2E, F), where different nutrients were used to achieve different doubling times (TYSON *et al*. 1979). In contrast, in other studies (GUO *et al*. 2004; BRAUER *et al*. 2008), which employed chemostats to alter the population doubling time independently of the limiting nutrient, there was no correlation between cell size and T_d_ or T_G1_ (Figure 2). Importantly, within these chemostat studies, cell size measurements were internally calibrated. Hence, the lack of any correlation between cell size and T_d_ or T_G1_ cannot be attributed to experimental variabilities of different studies incorporated in our meta-analysis of the literature. Lastly, the lack of a significant association between T_G1_ and cell size agrees with a genome-wide survey of single gene deletions (HOOSE *et al*. 2012), which found no pattern of correlation between cell size and the relative duration of the G1 phase. Hence, although cell size can be modulated by changes in nutrient composition in yeast ((TYSON *et al*. 1979; SOMA *et al*. 2014); and others), our data suggest that it is more likely that these are nutrient-specific effects, not causally linked to changes in cell proliferation rates.

### Cln3p, but not Whi5p, is required for the strong association between T_G1_ and T_d_

To understand how the relation between the length of the G1 phase and doubling time is established in budding yeast, we next examined the role of the Cln3p and Whi5p proteins, which regulate the G1/S transition in this organism. It has been proposed that dilution of Whi5p as cells get bigger in G1 is the key event controlling the timing of the G1/S transition (SCHMOLLER AND SKOTHEIM 2015; SCHMOLLER *et al*. 2015). Since in cells proliferating at different rates there is not a significant correlation with cell size (Figure 2, Table S1), it is not clear how the inhibitor dilution model would apply to the conditions we examine in this study. To our knowledge, the kinetics of cell cycle progression in cells lacking Cln3p or Whi5p have not been examined previously in steady-state cultures proliferating at different rates. To test the role of Cln3p and Whi5p in the relation between T_G1_ and T_d_, we examined the cell cycle profile of *cln3Δ* or *whi5Δ* cells in continuous, steady-state chemostat cultures. We measured T_G1_ in *cln3Δ* or *whi5Δ* cells from the DNA content of the cultures under glucose (0.08% ^W^/_v_) limitation and at different dilution rates (0.038 h^-1^ to 0.348 h^-1^; corresponding to T_d_ values between 18 h and 2 h, respectively). As expected, cells lacking Whi5p were very small (≈18 fL), and their size did not change significantly as a function of T_d_ (Figures 3E and S3D). Furthermore, the intercept of the linear fit between the log-transformed T_d_ and T_G1_ values of *whi5Δ* cells was significantly higher than the intercept of the linear fit of these parameters in wild-type cells (1.17 vs. 0.92; see Table 1), consistent with the shortened G1 phase of *whi5Δ* cells. The slope of the linear relation between T_d_ and T_G1_ in *whi5Δ* cells (Figure 3B) was similar to what we observed in the aggregate analysis of wild-type cells (Figure 1A, and Table 1; Sen-Adichie V statistic = 1.775, p-value = 0.183), albeit slightly smaller than the slope of wild type cells from a separate, independent experiment performed in this study (0.6723 in wild type (Fig. 3A) vs. 0.6166 in *whi5Δ* cells (Fig. 3B)). These data suggest that in different physiological states, and despite their shortened G1 phase and small size, *whi5Δ* cells nonetheless remain responsive to different environments, displaying minimal changes in their scaling of the expected proportional changes between T_d_ and T_G1_.

In contrast, cells lacking Cln3p had an abnormal behavior. In three independent experiments (Figures 3C, and S3B,C), T_G1_ did not even have a straightforward linear relation with T_d_ in cultures of cln3Δ cells. At shorter division times (T_d_ < 5 h), the T_d_ and T_G1_ values of cln3Δ cultures were related linearly, albeit with a higher slope (e.g., Figure S3C; slope = 0.7685). More importantly, in all three independent experiments, the linear relation breaks down at slower proliferating cln3Δ cultures (Figures 3C, and S3B,C). Even when combining all data points from the individual experiments for each strain, we found that at all doubling times tested the linear relation between T_d_ and T_G1_ remains strong for whi5Δ cells (τ > 0.8, based on Kendall’s non-parametric test). The same is true for cln3Δ cells at values of LnT_d_ < 1.5 (corresponding to T_d_ values < 4.5 h). In contrast, the linear relation between T_d_ and T_G1_ is significantly weaker (τ < 0.5) for slower proliferating cln3Δ cells (LnT_d_ > 1.5). These data suggest that Cln3p is more important than Whi5p for imposing the proper scaling relation of T_d_ ∝ T_G1_ in wild type cells.

### Cln3p levels are strongly and positively associated with cell proliferation rates

Given the important role of Cln3p in establishing the proper relation between T_d_ and T_G1_ (Figures 3 and S3), we sought to measure the levels of Cln3p and Whi5p as a function of T_d_. There are no reports of the steady-state levels of Cln3p or Whi5p in cell populations proliferating at different rates in chemostats.

To measure Whi5p and Cln3p levels from the same cells, we generated a strain that carries *WHI5-TAP* and *CLN3-13MYC* alleles, providing the only source of these gene products in the cells, expressed from their endogenous chromosomal locations (Figure 4A, see Materials and Methods). The expressed proteins were epitope-tagged, but otherwise un-mutated, wild type Whi5p-TAP and Cln3p-(Myc)_13_. These cells were then cultured in continuous, steady-state chemostat cultures under glucose (0.08% ^w^/_v_) or leucine (0.0015% ^w^/_v_) limitation. Although the *CLN3-13MYC* allele provides the means for reliable detection of otherwise wild type Cln3p, it is known to be slightly hypermorphic, stabilizing the Cln3p protein somewhat and shortening the G1 phase of the cell cycle (THORBURN *et al*. 2013). Indeed, the intercept of the linear fit between the log-transformed T_d_ and T_G1_ values of this strain (HB94; *WHI5-TAP, CLN3-13MYC*) was slightly higher than the intercept of the linear fit of these parameters in wild-type cells (0.97 vs. 0.92; see Table 1), consistent with a shortened G1 phase. The slope of the linear relation between the log-transformed values of T_d_ and T_G1_ was increased somewhat for these cells compared to the aggregate analysis of wild-type cells (0.68 vs. 0.6; see Table 1). Importantly, these cells still displayed a strong, linear, positive association between T_d_ and T_G1_ (τ=0.77; r=0.92; see Table 1), at all dilution rates we tested. Hence, we concluded that the relation between T_d_ and T_G1_ was only minimally affected in this strain, and we proceeded to quantify the levels of both Whi5p-TAP and Cln3p-(Myc)_13_, from separate chemostat experiments under glucose or leucine limitation, each run at ≥5 dilution rates (Figure 4; see Materials and Methods).

The levels of Whi5p-TAP were not increased in slower proliferating cells (Figure 4A, B). Previously, (LIU *et al*. 2015) reported that Whi5p abundance increases ≈3-fold in cells growing in poorer carbon sources, although this was not seen in a more recent study by (DORSEY *et al*. 2018). In any case, several variables were different between the (LIU *et al*. 2015) study and ours, which could account for the disagreement in the findings: First, different epitope-tagged alleles were used (*WHI5*-*tdTomato vs*. *WHI5-TAP*). Second, different detection methods were applied (fluorescence live cell imaging vs. immunoblots). Third, (LIU *et al*. 2015) use cells in the W303 background, which are larger than cells of the BY background we use here, possibly leading to differences in cell size regulation. Fourth, and most significantly, nutrient-specific effects could not be separated from growth rate-specific ones in (LIU *et al*. 2015). As we discussed earlier (see Figure 2), chemostats provide the only experimental approach for properly studying how rates of cell proliferation may affect a given output, separately from any effects unique to particular nutrients.

We observed a significant and disproportionate reduction of Cln3p-(Myc)_13_ levels in slower-proliferating cells (Figure 4A, B). With Cln3p-(Myc)_13_ levels normalized against the total cellular protein content, the fastest proliferating populations had >10-fold higher levels of Cln3p-(Myc)_13_ compared to the slowest proliferating cells (Figure 4A, B). Furthermore, because these estimates rely on the hypermorphic *CLN3-13MYC* allele that produces a slightly stabilized Cln3p protein (THORBURN *et al*. 2013), the dynamic range of Cln3p levels as a function of doubling time is likely even broader.

### A uORF in *CLN3* adjusts the levels of Cln3p at different cell proliferation rates

What is the mechanism that underpins the growth-dependent control of Cln3p abundance? We had predicted that a uORF in *CLN3* could inhibit its translational efficiency in poor media disproportionately (POLYMENIS AND SCHMIDT 1997). However, predictions of a growth-dependent role of the uORF had not been accompanied with measurements of Cln3p levels. A kinetic model of protein synthesis (LODISH 1974) forecasts that removing the uORF would de-repress synthesis of Cln3p in slowly dividing cells when the ribosome content of the cell is low. In contrast, removing the *CLN3* uORF would have minimal effects in cells that proliferate fast, when the ribosome content is high. To test this model, we introduced an A-315T substitution that mutates the start codon of the uORF in CLN3 without affecting CLN3 mRNA levels (POLYMENIS AND SCHMIDT 1997), in the strain that expresses otherwise wild type Whi5p-TAP and Cln3p-(Myc)_13_ (see Materials and Methods). Note also that in chemostat conditions very similar to the ones we used here, the levels of wild type *CLN3* mRNA do not change significantly as a function of growth rate (BRAUER *et al*. 2008).

The effects of the uORF were evident in slower proliferating cultures (T_d_ > 4h), where the dynamic range of Cln3p levels was much narrower (3-4 fold) in A-315T cells, and very different from the range of Cln3p levels (10-fold) in their wild type counterparts at these longer doubling times (p = 0.03648, based on the non-parametric Kolmogorov-Smirnov test), and indistinguishable from the range of Whi5p levels (Figure 4A, B and File S1). We note that although the range of Cln3p levels is narrower in slower proliferating *CLN3* uORF mutant cells (Figure 4), it is not flattened, arguing for additional mechanisms that could adjust the levels of Cln3p at different growth rates. Nonetheless, an independent piece of functional evidence further strengthened a growth-dependent role of the *CLN3* uORF. A hallmark phenotypic readout of gain-of-function *CLN3* alleles is a reduction in cell size (NASH *et al*. 1988). Cells that lack the *CLN3* uORF were smaller than their wild type counterparts were, and this effect was T_d_-dependent (see Figure 4C, D). Especially in leucine-limited cells, which displayed pronounced enlargement as they proliferated slower, removing the *CLN3* uORF reduced their size substantially (Figure 4D). These results are consistent with a de-repression of Cln3p synthesis upon removal of the *CLN3* uORF.

These data argue that translational control contributes to the disproportionate reduction of Cln3p levels as a function of T_d_. Note also that loss of Cln3p severely perturbs the linear relation between T_d_ and T_G1_ (Figures 3 and S3). In summary, our results underscore the critical role of the G1 cyclin Cln3p in the physiological coupling between growth and division.

## DISCUSSION

The scaling relations between G1 length and population time in yeast and human cells we report are significant for several reasons. First, if the duration of G1 is estimated, they allow predictions of proliferation rates, which could be useful in diverse settings, such as in tissues at an organismal level. Second, they serve as benchmarks against which the effects of genetic or other perturbations can be evaluated, as we demonstrated for Whi5p and Cln3p, two cell cycle regulators in yeast. Third, scaling relations of cellular physiology may ultimately point to general, physical mechanisms that organize life at the cellular level. In the next paragraphs, we discuss our findings in relation to current models of how cell division is controlled by cellular biosynthetic capacity, with emphasis on the roles of Whi5p and Cln3p.

### What is the context of this study in relation to others?

Our results pertain to cell cycle kinetics of steady-state cultures that proliferate at different rates, not to cell cycle adjustments immediately after nutrient shifts (TOKIWA *et al*. 1994; LEITAO AND KELLOGG 2017). We also did not examine G1 progression in a particular cell cycle, where small daughter cells will not initiate a new round of cell division until they reach a size characteristic of the culture medium (HARTWELL AND UNGER 1977; JOHNSTON *et al*. 1977). Hence the scaling of G1 duration between populations with different doubling times may not necessarily be controlled by the same mechanism that controls how G1 duration is regulated to maintain size homeostasis within a population of cells that proliferates at a given rate. This interpretation is consistent with the findings that cell size is not associated with rates of cell proliferation (Figure 2), at least in experimental settings of steady-state chemostat cultures, which separate nutrient-specific effects from the impact of different rates of cell proliferation. For example, note the very different sizes of cells in glucose vs. leucine-limited cultures at the same dilution rate (compare Figures 4C and 4D), offering yet another demonstration of how particular nutrients may affect cell size independently of any changes in rates of cell proliferation. Furthermore, in leucine-limited cultures, the cells were not only bigger than cells in glucose-limited chemostats at all dilution rates tested, but they also got even bigger as they divided slower (Figure 4). These observations argue against the widely-held assumptions that cells get smaller the slower they proliferate. Instead, they support the notion that nutrient effects on cell size may be particular to specific nutrients, and not associated with changes in rates of cell proliferation.

### How do our results mesh with models of G1 control?

As we noted above, our data were from cells dividing at different rates, which was not addressed in the Whi5p dilution model of (SCHMOLLER *et al*. 2015). Hence, the two studies are not directly comparable. The Whi5p dilution model, however, was constructed on the basis that while Whi5p levels were disproportionately lower than expected from cell growth, Cln3p levels were roughly constant and proportional to the increase in size from birth to START (SCHMOLLER *et al*. 2015). Given the disproportionate dependency of Cln3p levels on cell proliferation rates that we reported here and had predicted earlier (POLYMENIS AND SCHMIDT 1997), could the uORF-mediated translational control of *CLN3* affect Cln3p synthesis in the G1 phase from birth to START? We think this is unlikely because the uORF-mediated translational control we described operates when the concentration of active ribosomes in the cell changes (LODISH 1974), for example in poor vs. rich nutrients. Hence, while such translational control mechanisms provide excellent conduits to disproportionately alter gene expression and communicate growth-related inputs to downstream mRNA targets, to our knowledge, there is no report of cell cycle-dependent changes in ribosome content. Other mechanisms, not due to changes in the translational efficiency of *CLN3*, may contribute to significant, periodic changes in Cln3p abundance in G1, within a given cell cycle.

There are conflicting reports in the literature about whether Cln3p ‘cycles’ in the cell cycle. Cln3p is of such low abundance that cannot be properly measured in the single-cell microscopy methods of (SCHMOLLER *et al*. 2015), because mutant *CLN3* alleles had to be used, producing extremely stabilized and dysfunctional Cln3p protein that accumulates at very high, but non-physiological levels, so that it can be visible with microscopy. The initial report claiming that Cln3p-HA levels were constant in the cell cycle did not interrogate the early G1 phase (TYERS *et al*. 1993). In that report, although early G1, small (25 fL), elutriated daughter cells were collected, Cln3p levels were not measured until much later in G1 (at 35 fL, when by 40 fL 25% of the cells were already budded in that experiment; see Fig. 4 in (TYERS *et al*. 1993)). Based on that result, it had been assumed for decades that Cln3p levels were constant in the cell cycle. Recently, however, two independent studies by the Amon (THORBURN *et al*. 2013) and Kellogg (ZAPATA *et al*. 2014) labs, assayed elutriated synchronous cells carrying epitope-tagged, but otherwise wild type *CLN3* alleles. Both studies showed that Cln3p levels change >10-fold in G1. Cln3p is absent in early G1 cells, while it rises dramatically before START. We also used the same *CLN3-13MYC* allele to monitor Cln3p levels at different rates of cell proliferation (see Figure 4). The *CLN3-13MYC* allele is known to produce a slightly stabilized Cln3p protein (THORBURN *et al*. 2013). Note that, on the face of the slight stabilization of the Cln3p-(Myc)_13_, the dynamic range of Cln3p levels as a function of growth rate we report is likely even broader, not narrower. Hence, our conclusions are strengthened, not weakened by the slight stabilization of the Cln3-Myc we used. For the same reasons, the changes in Cln3p levels in G1 observed previously (THORBURN *et al*. 2013; ZAPATA *et al*. 2014) are likely even greater than indicated in these reports.

Overall, although a 2-fold dilution of Whi5p is observed in G1 (SCHMOLLER *et al*. 2015), the changes in Cln3p levels are likely more pronounced (THORBURN *et al*. 2013; ZAPATA *et al*. 2014), through transcriptional (ZAPATA *et al*. 2014), or other mechanisms. In this context, it is perhaps unsurprising that we found Cln3p to be more important than Whi5p in the relation between T_d_ and T_G1_. It is important to stress, however, that any changes of Cln3p levels in G1 do not affect the key aspect of the inhibitor dilution model, namely that Whi5p levels are reduced by cell growth. Hence, we may be dealing with a more complex, ‘mixed’ model of inhibitor dilution and activator accumulation. It is possible that the levels of additional proteins may behave analogously to Cln3p and Whi5p, contributing to a broader network of factors whose antagonistic relations control the timing of initiation of cell division. Regardless of the identity of those proteins, in yeast and other models, the fundamental relation between T_G1_ and T_d_ we describe in this report will serve as a useful metric to evaluate the role of these protein(s) in the control of cell division by growth inputs.

## ACKNOWLEDGEMENTS

### Author Contributions

Methodology, H.M.B., M.P.; Formal Analysis, H.M.B., M.P.; Investigation, H.M.B., M.C., I.P.E.P., A.O.P., M.P.; Resources, M.P.; Data Curation, H.M.B., M.P.; Writing – Original Draft, M.P., Writing – Review and Editing, H.M.B., M.P.; Visualization, H.M.B., M.P.; Supervision, H.M.B., M.P., Funding Acquisition, M.P.

This work was supported from the NIH (grant GM123139 to M.P.). M.C. was supported by a National Science Foundation REU award (DBI-1358941). The authors declare no conflicts of interest.

